# Identification of Circular Dorsal Ruffles as Signal Platforms for the AKT pathways in Glomerular Podocytes

**DOI:** 10.1101/2022.06.22.497178

**Authors:** Rui Hua, Mauricio Torres, Jinzi Wei, Xiaowei Sun, Li Wang, Ken Inoki, Sei Yoshida

## Abstract

Circular dorsal ruffles (CDRs) are rounded membrane ruffles induced by growth factors to function as precursors of the large-scale endocytosis called macropinocytosis. In cell line systems, CDR/macropinocytosis regulate the AKT-mTORC1 pathway, a canonical growth factor signaling. However, it is not known if this mechanism occurs in tissues. Here, utilizing ultra-high-resolution scanning electron microscopy, we report that CDRs are expressed in glomerular podocytes *ex vivo* and *in vivo*. Biochemical and imaging analysis revealed that AKT phosphorylation is localized to CDRs upstream of mTORC1 activation in podocyte cell line and isolated glomeruli, indicating that CDRs function as signal platforms for AKT-mTORC1 pathway in podocytes to maintain the kidney function. Because mTORC1 has critical roles in the podocyte metabolism and the aberrant activation of mTORC1 triggers podocytopathies, these results suggest that targeting CDR formation would be a potential therapeutic approach for the diseases.

## INTRODUCTION

Circular dorsal ruffles (CDRs) are crater-shaped membrane ruffles that appear on cell surfaces stimulated by growth factors, such as platelet-derived growth factor (PDGF), hepatocyte growth factor (HGF), insulin, and epidermal growth factor (EGF) (Hoon *et al*, 2012; Itoh & Hasegawa, 2013). The molecular mechanism and cellular functions of CDRs have been studied in the past decade (Hoon *et al*., 2012; Itoh & Hasegawa, 2013); our research group and others have found that CDRs transform into macropinocytosis, a large-scale solute uptake endocytosis, in fibroblasts and glioblastoma cells (Bernitt *et al*, 2017; Dharmawardhane *et al*, 2000; Gu *et al*, 2011; Legg *et al*, 2007; Yoshida *et al*, 2018b; Zdzalik-Bielecka *et al*, 2021). Live-cell imaging showed that after GF stimulation, membrane protrusions were evoked from the cell surface to become CDRs and they gradually shrunk towards the center of the structure and generated macropinosomes (Yoshida *et al*., 2018b; Zdzalik-Bielecka *et al*., 2021).

GF stimulation activates several canonical signaling pathways, such as the cascade including phosphoinositide 3-kinase (PI3K), mechanistic target-of-rapamycin complex-2 (mTORC2), AKT, and mechanistic target-of-rapamycin complex-1 (mTORC1) (Fu & Hall, 2020; Hoxhaj & Manning, 2020; Liu & Sabatini, 2020; Yoshida *et al*, 2018a). After GF stimulation, PI3K is activated and generates phosphatidylinositol (3,4,5)-triphosphate (PIP3) at the plasma membrane. The protein kinase AKT contains a pleckstrin homology (PH) domain that interacts with PIP3. Thus, once PIP3 is generated, AKT is recruited to the plasma membrane, where the protein is phosphorylated and activated by two distinct kinases, PDK1 and mTORC2. Phospho-AKT functions upstream of mTORC1, which has a key role in cell growth and differentiation.

Recent studies have shown that CDR and macropinocytosis modulate the PI3K/AKT/mTORC1 signaling pathway (Stow *et al*, 2020; Swanson & Yoshida, 2019; Yoshida *et al*., 2018a). Imaging analysis showed that PIP3 accumulates at macropinocytic cups in macrophages, resulting in the local recruitment of AKT (Pacitto *et al*, 2017; Wall *et al*, 2017; Yoshida *et al*, 2009). The inhibition of macropinocytosis was found to block AKT phosphorylation (pAKT) and mTORC1 activation in macrophages (Pacitto *et al*., 2017; Yoshida *et al*, 2015). A number of studies have shown that the expression of oncogenic Ras induces both macropinocytosis and mTORC1 activation (Bar-Sagi & Feramisco, 1986; Egami *et al*, 2014; Inoki *et al*, 2003; Mendoza *et al*, 2011), and the macropinocytosis-specific inhibitor EIPA is reported to diminish Ras-induced mTORC1 activation (Palm *et al*, 2015). NUMB protein negatively regulates CDR formation; depletion of NUMB increases HGF-induced pAKT (Zobel *et al*, 2018). Moreover, we recently described PIP3 generation and AKT recruitment to CDRs, and inhibition of CDR formation was shown to attenuate EGF-induced AKT phosphorylation (Sun *et al*, 2022; Yoshida *et al*., 2018b).

Podocytes are highly differentiated renal epithelial cells that cover the glomerular capillaries as an essential component of the kidney filtration barrier (Assady *et al*, 2017; Garg, 2018; Kopp *et al*, 2020), and CDR formation and macropinocytosis have been shown to occur in these cells (Bollee *et al*, 2011; Chung *et al*, 2015; Schell *et al*, 2013). Heparin-binding epidermal growth factor-like growth factor (HB-EGF) was shown to induce CDR formation in cultured mouse podocytes (Bollee *et al*., 2011). EGF treatment generates CDRs in primary cultured podocytes (Schell *et al*., 2013). Staining of mouse tissue revealed that podocytes have active macropinosomes *in vivo* (Chung *et al*., 2015). However, the cellular function of CDRs and the interaction between CDRs and macropinocytosis in podocytes remain unclear, and microscopic detection of CDRs *in vivo* has not been reported. Thus, the physiological relevance of CDRs in podocytes should be investigated *in vivo* as well as *ex vivo*.

We hypothesized that podocyte CDRs precede macropinocytosis and regulate the AKT/mTORC1 pathway to maintain cell metabolism as a critical part of kidney function. In the current study, we tested this idea using different techniques to identify CDRs on podocytes *in vivo* and *ex vivo* and investigated their role in the GF-induced AKT/mTORC1 pathway. High-resolution scanning electron microscopy (SEM) of mouse kidney tissue showed that glomerular podocytes displayed CDR-like structures *in vivo*. Moreover, both EGF and PDGF induced CDRs in the mouse podocyte cell line, MPC5. Inhibition of CDR formation significantly mitigated GF-induced AKT phosphorylation and attenuated mTORC1 activation in cells. Confocal microscopy showed that PI3K/mTORC2/AKT signaling components localized to GF-induced CDRs. Importantly, we utilized isolated mouse glomeruli for *ex vivo* experiments and found that glomerular podocytes express CDRs as macropinocytic cups, regulating the AKT pathway. Thus, with these results, we have named the cellular function of CDRs *ex vivo* and *in vivo*, and we propose that CDRs are signaling platforms for growth factor-induced AKT pathways in podocytes as a critical part of the kidney filtration function.

## RESULTS AND DISCUSSION

### Podocytes express CDRs as a precursor of macropinocytosis

Ultra-high-resolution SEM of mouse glomeruli was used to further characterize the morphology of podocytes. Unexpectedly, we found crater-shaped membrane ruffles on the surfaces of glomeruli (**Fig. 1, arrows**). Higher-magnification images show rounded membrane ruffles (**Fig. 1, enlarged images**) protruding from the podocyte cell body, which is the center of podocytes, and an elongated unique cellular structure called the primary/foot process. We identified a total of 21 glomeruli from five mice and found that 25.38% of 528 podocytes induced these structures.

**Figure 1.**
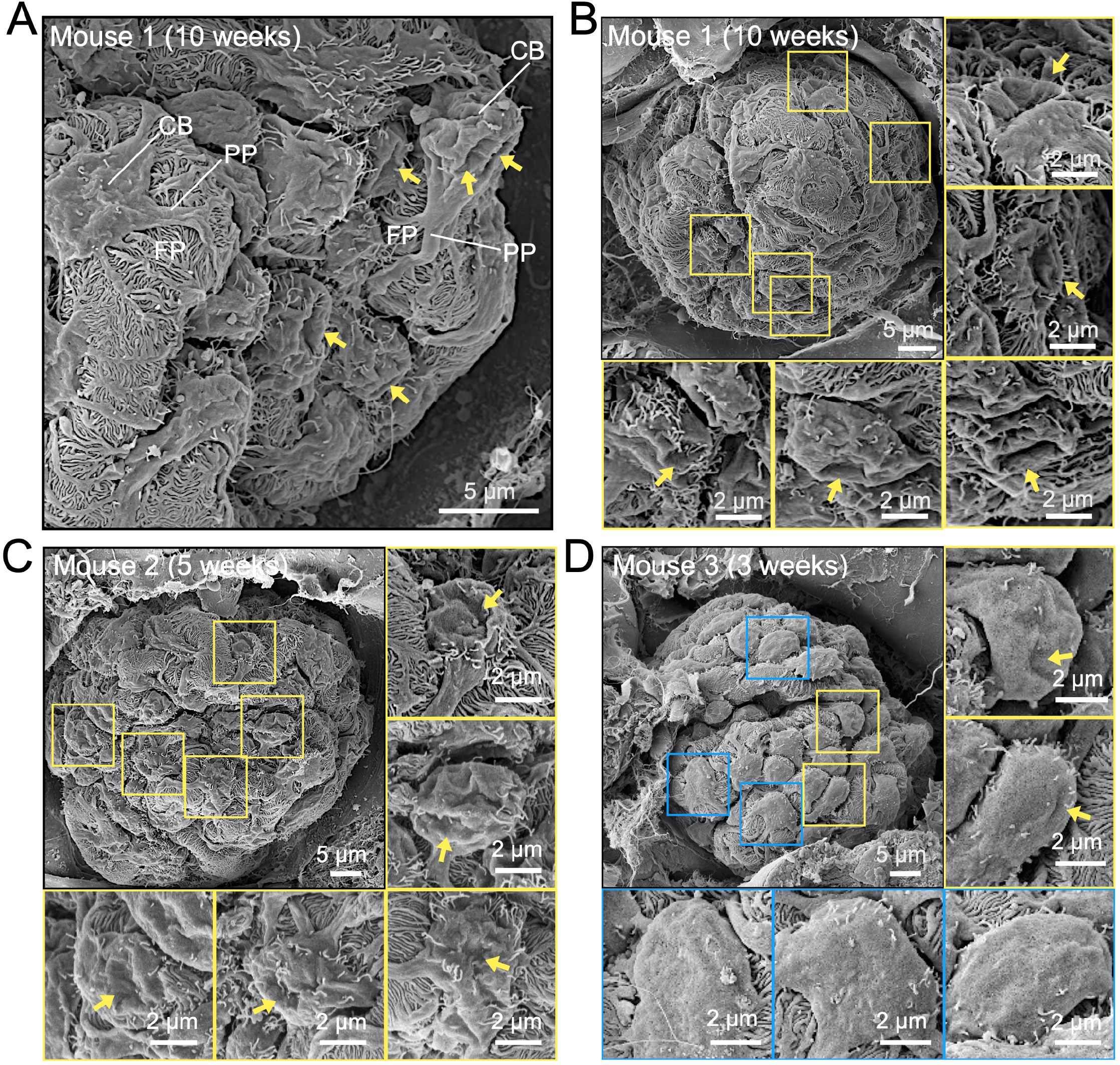
Representative high-resolution SEM images of mouse glomeruli showing circular dorsal ruffle (CDR)-like structures in podocytes. In total, 21 glomeruli from 5 mice were examined using SEM. Arrows indicate CDR-like structures; squares indicate podocytes with (yellow) or without (blue) CDR-like structures. CDR-like structures were induced in 25.38 % of the podocytes (n=528). CB, the cell body of podocytes; PP, primary process; FP, foot process. Scale bars: 5 μm and 2 μm (enlarged insets). Glomeruli are from mice aged 10 weeks (**A and B**), 5 weeks (**C**), and 3 weeks (**D**).

Membrane ruffles can be induced by growth factors (Hoon *et al*., 2012; Itoh & Hasegawa, 2013; Kay, 2021; Swanson, 2008). Thus, we analyzed the EGF-stimulated mouse podocyte cell line, MPC5. Actin was stained to identify CDRs (Hoon *et al*., 2012; Itoh & Hasegawa, 2013; Sun *et al*., 2022), and antibodies against WT1 were used as a marker of podocytes (Guo *et al*, 2002; Inoki *et al*, 2011; Yoshida *et al*, 2021). The results showed that actin formed ring structures covering the central part of the cells (**Fig. 2A, EGF, arrows**), suggesting that EGF induced CDRs in these cells. Quantification analysis from 30 independent experiments confirmed that EGF induced CDRs in MPC5 at an efficiency of 30.17% in 63,606 cells analyzed. We also found that PDGF induced CDRs in MPC5 (10.91% of 65,287 cells from 30 independent experiments) (**Fig. 2A, PDGF, arrows**). Interestingly, podocytes also induced CDRs—although rarely—under culture conditions (**Fig. 2A, Culture condition, arrows**). Recruitment of Rab5a (Lanzetti *et al*, 2004; Palamidessi *et al*, 2008), SH3YL1 (Hasegawa *et al*, 2011), and cortactin (Cortesio *et al*, 2010) were used as the CDR marker (Sun *et al*., 2022). We also found that these proteins were expressed at GF-induced CDRs (**Fig. 2B and Fig. S1A and B**). Thus, we concluded that GF treatment induces CDRs in podocytes.

**Figure 2.**
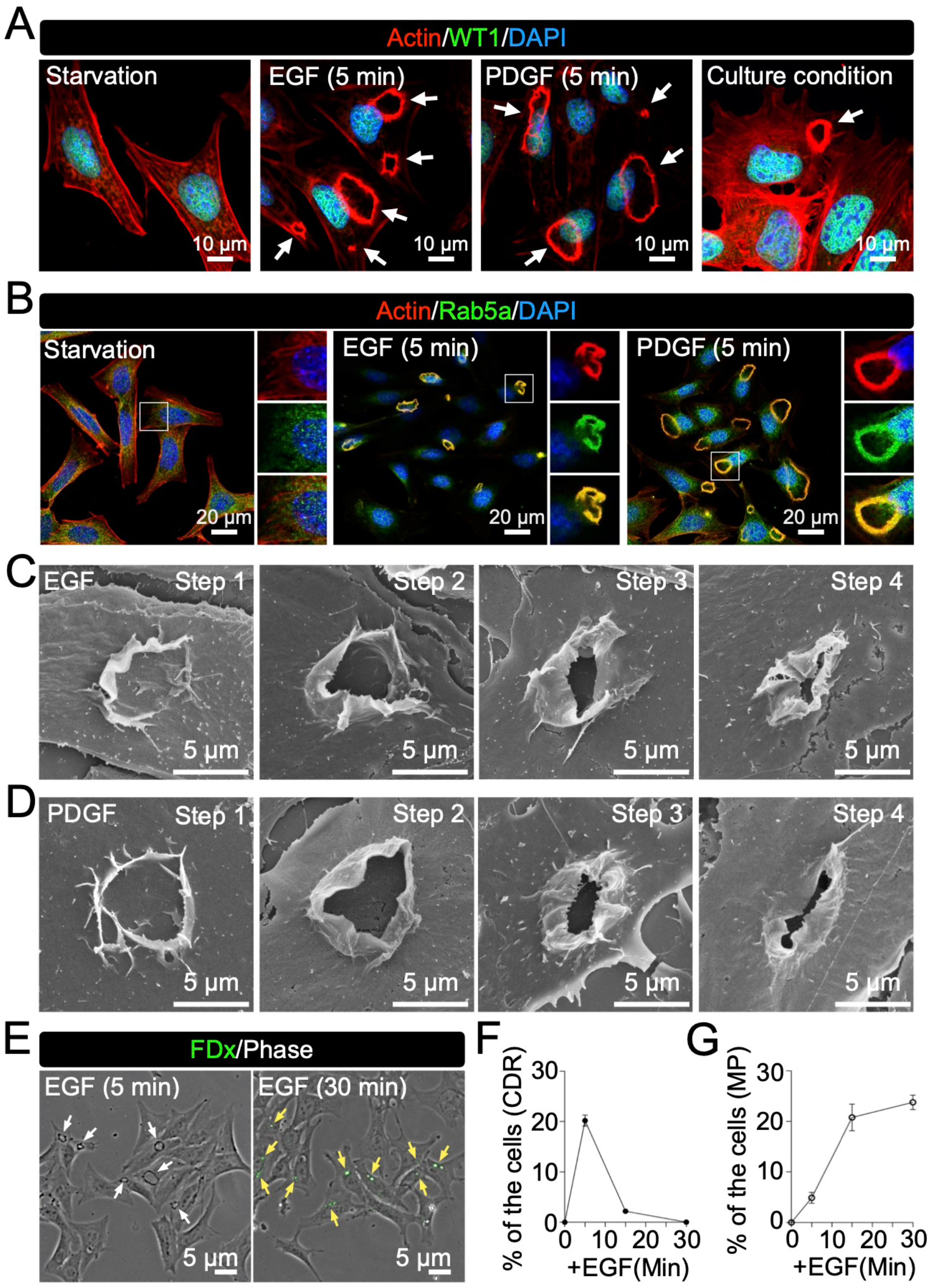
EGF and PDGF induce CDRs in mouse podocyte cell line MPC5. (**A and B**) Representative confocal images of GF-treated MPC5. Treatment with epidermal growth factor (EGF) and platelet-derived growth factor (PDGF) for 5 min induced CDRs (white arrows) identified by actin staining (red). Transcription factor WT1 (green) was used as a podocyte marker (**A**). Rab5a was used as a CDR marker (**B**). (**C and D**) Representative high-resolution SEM images of CDRs induced by EGF (**C**) or PDGF (**D**) show four different morphological patterns. (**E**) Imaging analysis identified CDRs (white arrows in 5 min) and macropinosomes identified by FDx70 (yellow arrows in 30 min) in MPC5 after EGF treatment. (**F and G**) Quantification analysis from three independent time-course experiments revealed that while CDRs disappeared (**F**) macropinosomes were generated (**G**) during the observation. Scale bars: 10 μm (**A**), 20 μm (**B**), and 5 μm (**C-E**).

The results also revealed different sizes of CDRs in the cells after stimulation (**Fig. 2A, arrows**), suggesting that CDRs shrink to the center and form macropinosomes, as observed in mouse embryonic fibroblasts (MEFs) (Yoshida *et al*., 2018b). Ultra-high-resolution SEM was used to depict the structure of the CDRs in MPC5. We identified at least four different morphological steps:1) circular protrusions were evoked from the surface of the cell, 2) the structure was elongated in the vertical direction to form wall-like membrane ruffles, 3) the membrane ruffles were waved and bent to the inside at the edge, and 4) the top open area was closed by a fusion of the ruffles (**Fig. 2C and D**), suggesting that the CDRs function as macropinocytic cups. To evaluate the ingestion activity of CDR via macropinocytosis, we used FDx70 as a probe for the ingestion of extracellular solutes. Phase-contrast images clearly showed CDR structures 5 min after EGF stimulation (**Fig. 2E, white arrows**) and FDx70 fluorescence appeared as circular structures in the cells after 30 min (**Fig. 2E, yellow arrows)**. In fact, the time-course experiment revealed that, whereas the number of induced CDRs decreased, the signal intensity of FDx70 increased during the observation period (**Fig. 2F and G**), suggesting that the CDRs changed to macropinosomes and ingested FDx70. These data indicate that GF-induced CDRs form macropinosomes in podocytes, which ingest extracellular solutes.

### CDRs are required for the AKT-mTORC1 pathway in podocytes

EGF and PDGF activate the AKT/mTORC1 pathway in different cell lines (Chen *et al*, 2016; Hoxhaj & Manning, 2020; Ying *et al*, 2017). Accordingly, we confirmed that GF induced AKT phosphorylation (pAKT) in MPC5 within 1 min of stimulation (**Fig. S1C and D**). As an output of mTORC1 activation, S6K phosphorylation (pS6K) was also evident. Following stimulation with both EGF and PDGF, pS6K expression was induced for 5 min (**Fig. S1C and D**), thus supporting the hypothesis that AKT acts upstream of mTORC1. ERK phosphorylation (pERK), another downstream signaling molecule of GF stimulation, was observed at 1 min (**Fig. S1C and D**). To test the role of CDRs in growth factor signaling, we used the macropinocytosis-specific inhibitor EIPA (Koivusalo *et al*, 2010). EIPA treatment diminished GF-induced pAKT and pS6K (**Fig. 3A and B**) and completely blocked GF-induced CDR formation (**Fig. 3G-J**), suggesting that CDRs are required for the AKT-mTORC1 pathway.

**Figure 3.**
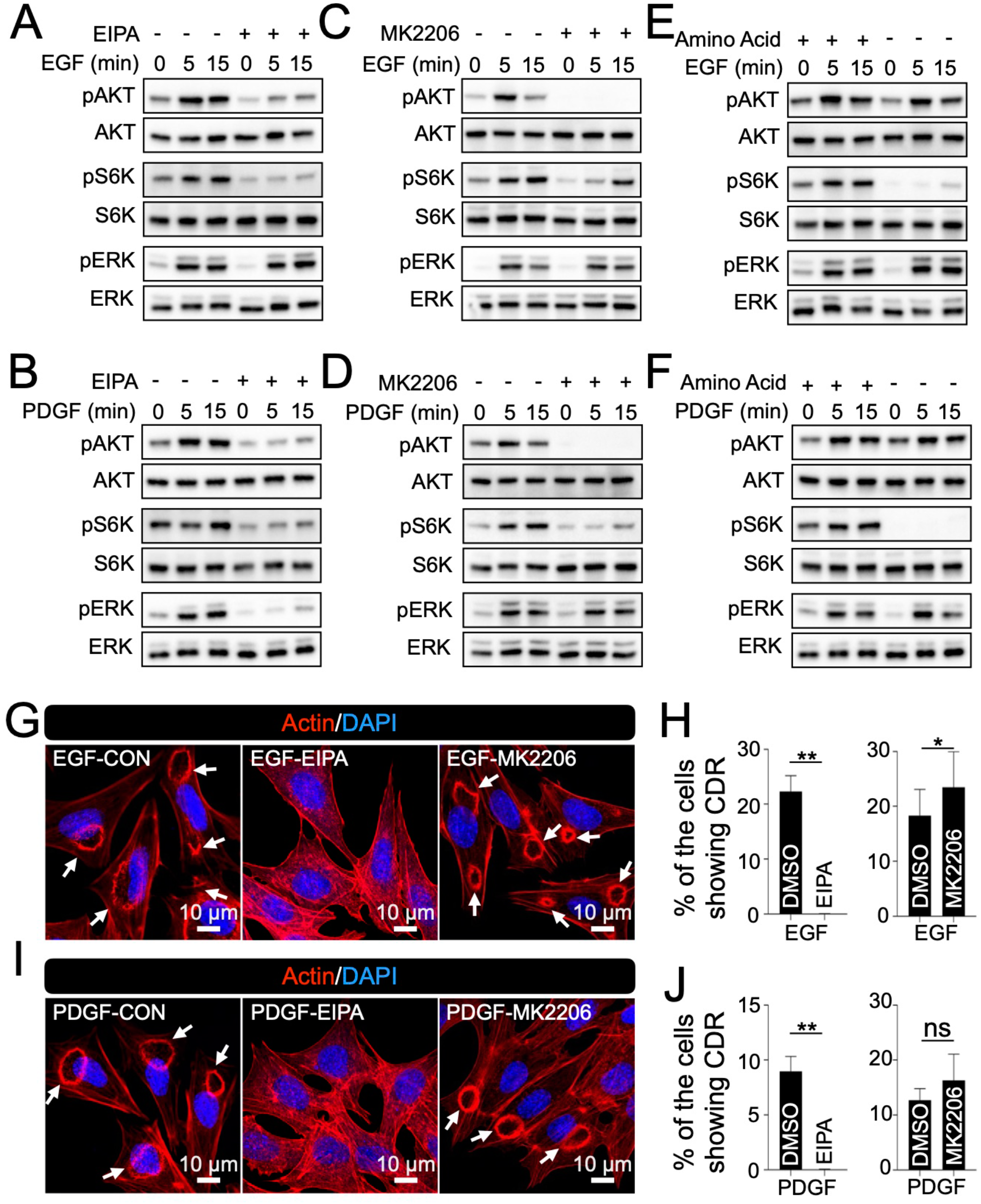
CDRs regulate GF-induced AKT/mTORC1 pathway in MPC5. Inhibition of CDRs attenuated pAKT and pS6K. (**A and B**) Western blot analysis of AKT, S6K, and ERK signals after EGF (**A**) or PDGF (**B**) stimulation with/without macropinocytosis inhibitor EIPA. (**C and D**) Western blot analysis of AKT, S6K, and ERK signals after EGF (**C**) or PDGF (**D**) stimulation with/without AKT inhibitor MK2206. (**E and F**) Western blot analysis of AKT, S6K, and ERK signals after EGF (**E**) or PDGF (**F**) stimulation with/without amino acids. (**G and I**) Representative confocal images of MPC5 stimulated by EGF (**G**) or PDGF (**I**) with/without EIPA or MK2206. Arrows indicate CDRs. Scale bar: 10 μm. (**H and J**) Results of CDR assays from five independent experiments showed that EIPA treatment blocked GF-induced CDRs, but MK2206 did not. **: p<0.01, *: p<0.05. Two-tailed paired Student t-test was applied to compare the two groups.

The actin polymerization protein N-WASP has been observed at CDRs, and the N-WASP inhibitor, wiskostatin, is known to block CDRs (Legg *et al*., 2007; Schell *et al*., 2013). We also found that the protein was located at CDRs (**Fig. S1E)** and that wiskostatin blocked GF-induced CDRs in MPC5 cells (**Fig. S1F and G)**. Biochemical analysis showed that wiskostatin treatment blocked GF-induced pAKT and pS6K, but not pERK (**Fig. S1H and I**), suggesting that CDRs regulate the AKT/mTORC1 pathway. When we used the AKT-specific inhibitor MK2206, we found that, although pAKT was completely blocked (**Fig. 3C and D**), CDRs were still induced by GFs (**Fig. 3 G-J**). Thus, AKT was not upstream of CDR formation. It has been shown that extracellular amino acids ingested by macropinocytosis activate mTORC1 in growth factor signaling (Yoshida *et al*., 2015). We found that GF treatment-induced pAKT, but not pS6K, under amino acid starvation conditions (**Fig. 3E and F**), suggesting that ingestion of extracellular amino acids is necessary for mTORC1 activation in podocytes. Based on these results, we concluded that CDRs in podocytes work upstream of the GF-induced AKT/mTORC1 pathway via two mechanisms: 1) CDRs function as a signaling platform for AKT activation, and 2) CDR ingests extracellular nutrients via macropinocytosis to activate mTORC1.

### The PI3K/mTORC2/AKT pathway is activated at CDRs

AKT and mTORC2 are recruited to the plasma membrane by interacting with PIP3, which is generated by PI3K. Our confocal microscopy observations revealed that the PI3K catalytic subunits p110α and p110β isoforms were recruited to the CDRs (**Fig. 4A** and **Fig. S2A and B**). Thus, we tested whether PIP3 is generated at CDRs by utilizing GFP-tagged BtK-PH, an established probe protein to localize PIP3 (Varnai *et al*, 1999; Yoshida *et al*., 2009). Confocal microscopy showed strong GFP-BtkPH signals at CDRs in GF-stimulated cells expressing probe proteins (**Figs. 4B and S2C**). mSin1 is a key component of mTORC2 (Fu & Hall, 2020). We also utilized GFP-AKT and GFP-mSin1 and found that these proteins were recruited to CDRs (**Figs. 4C-D and S2C**). These results suggest that CDRs are used as signaling platforms for the PI3K/mTORC2/AKT pathway.

**Figure 4.**
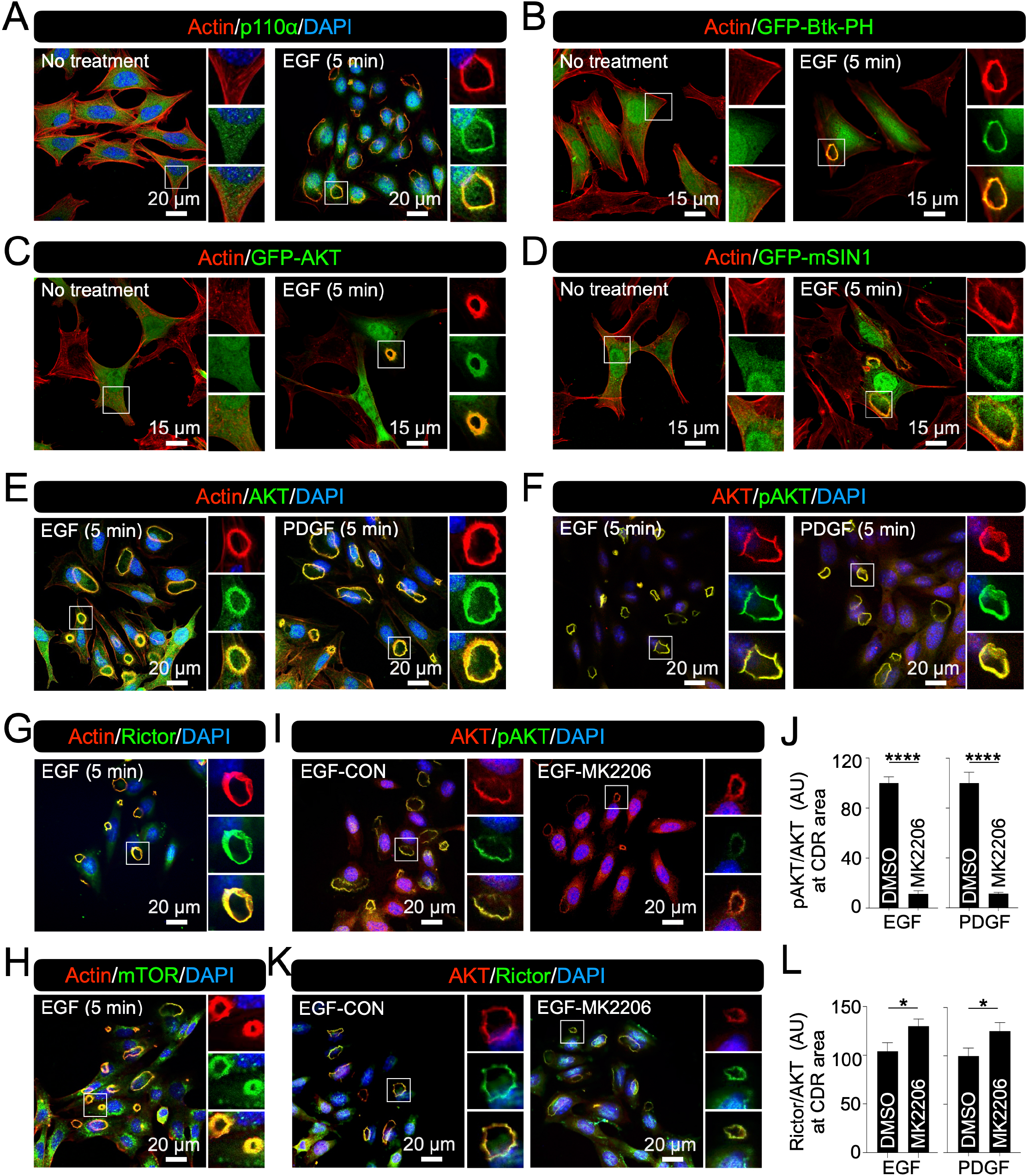
PI3K/mTORC2/AKT pathway is activated at CDRs in MPC5. (**A**) Representative confocal images of actin (red) and p110α (green) with and without EGF stimulation. (**B-D**) Representative confocal images of actin (red) and GFP-Btk-PH (green in **B**), GFP-AKT (green in **C**), GFP-mSIN1 (green in **D**) with/without EGF stimulation. (**E**) Representative confocal images of actin (red) and AKT (green) with EGF or PDGF stimulation. (**F**) Representative confocal image of AKT (red) and pAKT (green) with EGF or PDGF stimulation. (**G and H**) Representative confocal image of actin (red)/rictor (green) (**G**) and actin (red)/mTOR (green) (**H**) after EGF stimulation. (**I and K**) Representative confocal image of AKT/pAKT **(I)** and AKT/rictor (**K**) after EGF stimulation with and without MK2206. (**J and L**) Quantification of the ratio of pAKT/AKT (**J**) and rictor/AKT (**L**). More than 12 images from two independent experiments were analyzed. ****: p<0.0001, *: p<0.05. Two-tailed unpaired Student t-test was used for statistics. Scale bars: 20 μm (**A, E-I, and K**), and 15 μm (**B-D**).

To test whether AKT is phosphorylated by mTORC2 at CDRs, we performed several immunofluorescence (IF) staining. Similar to the results of the overexpression experiment, the endogenous AKT was found to be localized at CDRs (**Figs. 4E and S2D**). The staining patterns of AKT and pAKT (473) at CDRs completely matched (**Figs. 4F and S2D**), suggesting that AKT is phosphorylated at the CDRs by mTORC2. Rictor and mTOR are components of mTORC2 (Fu & Hall, 2020). The IF staining also revealed that both rictor (**Figs. 4G and S2E**), and mTOR (**Figs. 4H and S2F**) were localized at the CDRs. IF detection of LAMP, a marker of late endosomes, did not reveal the localization of LAMP to the CDR (**Fig. S2G**). Thus, LAMP is useful as a negative control to ensure that IF staining identifies specific signals at CDRs.

Western blot analysis showed that the AKT inhibitor MK2206 completely blocked GF-induced AKT phosphorylation (473) (**Fig. 3C and D**). Accordingly, once we treated cells with the inhibitor, the signal intensity of pAKT (473) at CDRs was significantly weaker compared to no treatment, whereas the intensities of AKT at CDRs in both cases were similar (**Figs. 4I-J and S2H**). In contrast, rictor was retained at the CDRs of the MK2206-treated cells (**Figs. 4K-L and S2I**), supporting the hypothesis that pAKT(473) is induced by mTORC2 at CDRs. Thus, these results strongly suggest that PIP3 generated at the CDRs recruits both AKT and mTORC2, leading to CDR-dependent AKT phosphorylation.

### Podocytes induce CDRs to modulate the AKT-mTORC1 pathway at the surface of the glomerulus

Although the results of our *in vivo* experiments clearly showed that glomerular podocytes induce CDRs (**Fig. 1**), the limitations of this approach meant that we could not identify the particular cellular function. Cell line experiments indicated that CDRs regulate the AKT pathway in podocytes (**Figs. 2-4**); however, the physiological relevance was not clear. To confirm these complementary results, we performed *ex vivo* experiments using isolated mouse glomeruli. Mouse glomeruli were isolated and treated with EGF according to the established method (Wang *et al*, 2019). As expected, ultra-high-resolution SEM revealed that there were CDRs expressed in podocytes on the surface of the glomerulus (**Fig. 5A, arrows**). As in the cell lines, CDRs were observed on the surface of the glomeruli, even without EGF treatment (**Fig. S3A, arrows**), suggesting that glomerular podocytes continuously expressed CDRs.

**Figure 5.**
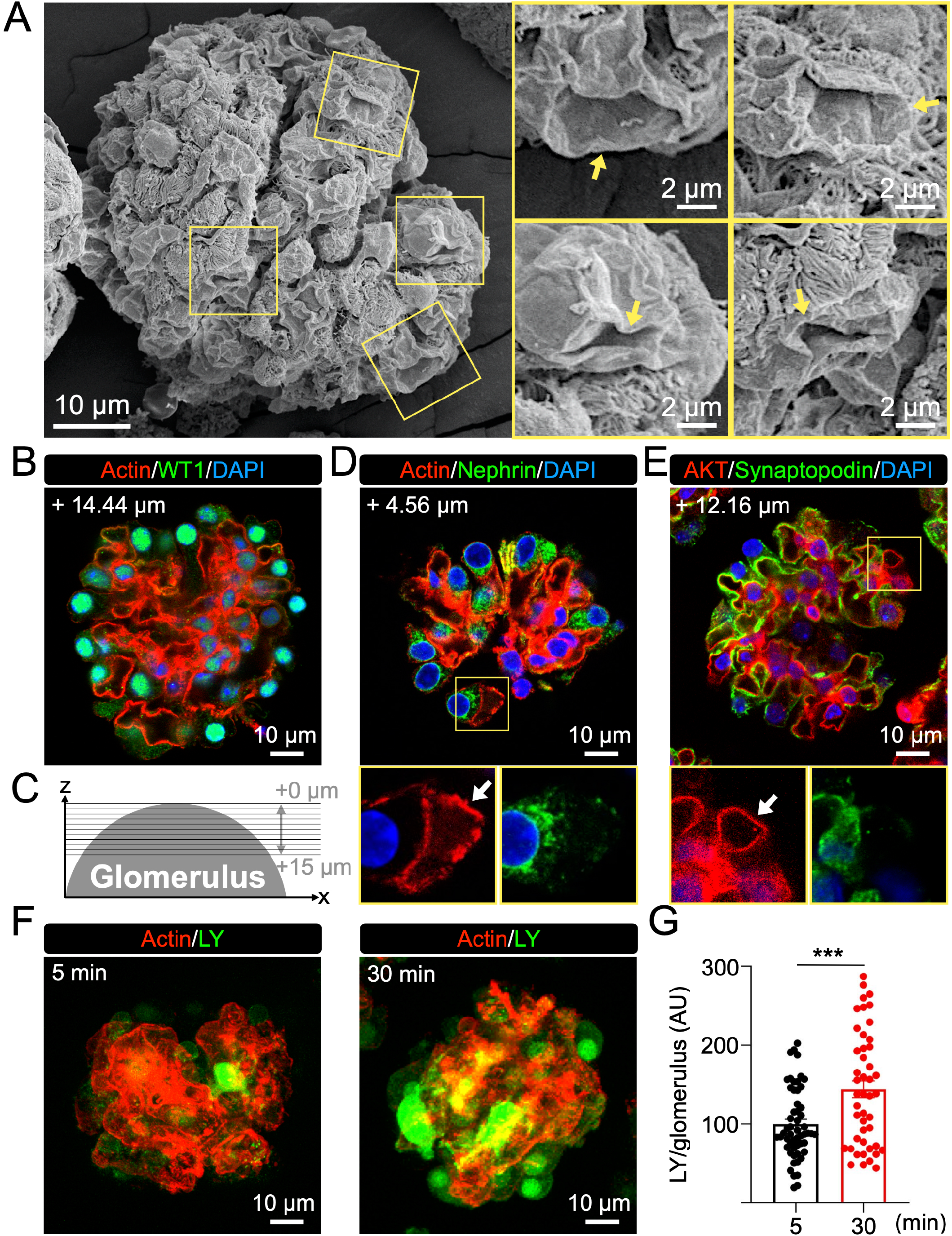
*Ex vivo* experiments identified AKT recruitment to the CDRs expressed in podocytes on isolated glomeruli. (**A**) Representative high-resolution SEM images of isolated glomerulus showing CDR-like structures (yellow squares). Isolated glomeruli were treated with EGF and fixed for SEM observation. Structures that resemble morphological steps of CDR formation in MPC5 were identified with high magnification (arrows). Scale bars: 10 μm and 2 μm (enlarged insets). (**B**) Representative confocal images of actin (red) and WT1 (green) in the isolated glomerulus. Podocytes identified by WT1 were observed at the edge of the glomerulus. Scale bar: 10 μm. (**C**) Schematic of a method to detect signals from an isolated glomerulus. Images of a glomerulus were scanned according to the Z-axis from the top as +0 μm each 0.76-μm interval until the +15 μm section, as illustrated. (**D**) Representative confocal image of actin (red) and nephrin (green) in an isolated glomerulus. CDR was identified as an actin-positive/nephrin-negative ring (arrow). Scale bar: 10 μm. (**E**) Representative confocal image of AKT (red) and synaptopodin (green) in an isolated glomerulus. AKT was observed at a CDR (arrow). Scale bar: 10 μm. The numbers shown in the images indicate the z-axis location of each section (**B, D, and E**). (**F**) Representative XY-projection images of actin (red) and LY (green) in an isolated glomerulus. Isolated glomeruli were treated with LY for 5 or 30 min. Scale bar: 10 μm. (**G**) Comparison of LY signal inside of isolated glomeruli between 5- and 30-min treatments. ***: p<0.001. Two-tailed unpaired Student t-test was used for statistics (n>40).

Utilizing isolated glomeruli allowed us to identify the location of each podocyte in three-dimensional images. After IF staining, we scanned the structure every 1-2 μm along the z-axis from the upper plane to 15 µm from the top of the shape (**Fig. 5C**). As expected, we observed WT1 positive cells at the edges of the isolated glomeruli (**Fig. 5B**), indicating that the cells observed at the edges of the structure were podocytes. Because nephrin and synaptopodin are expressed in the plasma membranes of podocytes for cell-cell interactions called slit diaphragms (Assady *et al*., 2017; Garg, 2018; Liu *et al*, 2001; Mundel *et al*, 1997a), they can be used to identify the edges of podocytes (Godel *et al*, 2011; Inoki *et al*., 2011; Puelles *et al*, 2019; Yoshida *et al*., 2021). Actin and nephrin double staining revealed actin-positive but nephrin-negative structures in cells (**Figs. 5D and S3B**), indicating that with this staining we can identify actin circles at the center of the cells. Moreover, we observed AKT-positive but synaptopodin-negative structures in the cells, suggesting that AKT was recruited to the actin ring (**Figs. 5E and S3C**). These results strongly suggest that AKT is recruited to the induced CDRs in glomerular podocytes *ex vivo*.

To test if these CDRs precede macropinocytosis, we used the fluid-phase probe Lucifer yellow (LY) as a representative extracellular molecule (Berthiaume *et al*, 1995; Yoshida *et al*., 2015), and the signal inside the glomerular podocytes was monitored in order to identify macropinosomes. Isolated glomeruli were treated with LY and incubated for 5 or 30 min. Imaging analysis revealed that the LY signal at 30 min was stronger than at 5 min (**Fig. 5F and G**). These results strongly suggest that CDR functions as a precursor of micropinocytosis at the surface of isolated glomeruli to ingest extracellular solutes.

### Proposed model: CDRs function as signal plat forms for the AKT pathways in glomerular podocytes

We identified CDRs in glomerular podocytes, both *in vivo* and *ex vivo* (**Figs. 1, 5A, and S3A**). Ultrahigh-resolution SEM observations revealed a morphological transition from CDRs to macropinosomes *in vitro* (**Fig. 2C and D**), and *ex vivo* (**Figs. 5A and S3A**). The results of FDx70 and LY assays confirmed that CDRs underwent macropinocytosis *in vitro* (**Fig. 2E-G**), and *ex vivo* (**Fig. 5F-G**), respectively. Confocal microscopy showed recruitment of GFP-Btk-PH, GFP-AKT, and GFP-mSin1 to CDRs (**Figs. 4B-D and S2C**), suggesting the recruitment of AKT and mTORC2 to the plasma membrane via their interaction with PIP3. IF staining for endogenous proteins localized to AKT, pAKT, mTOR, rictor, p110α, and p110β in the CDRs (**Figs. 4A-B, 4E-H, S2A-B, and S2D-F**). Moreover, the results of western blot analysis showed that the inhibition of CDRs blocked pAKT (**Figs. 3A-B and S1H-I**). Thus, we anticipate that GF-induced CDRs may function as signaling platforms for the PI3K/AKT pathway and contribute to the growth, development, and viability of podocytes.

Additionally, the inhibition of CDRs attenuated S6K phosphorylation (pS6K) (**Figs. 3A-B and S1H-I**), a well-known mTORC1 substrate. Previous studies have shown that macropinocytosis ingests extracellular nutrients and delivers extracellular amino acids to lysosomes, resulting in mTORC1 activation (Palm *et al*., 2015; Swanson & Yoshida, 2019; Yoshida *et al*., 2015). In fact, our data showed that EGF induced pAKT, but not pS6K, under amino acid starvation conditions *in vitro* (**Fig. 3E-F**). Thus, macropinosomes formed from CDRs would ingest extracellular nutrients and transport them to lysosomes to activate mTORC1. Here, we hypothesized that CDRs support mTORC1 activity via two mechanisms:

1. CDRs function as a signal platform for the PI3K-AKT pathway upstream of mTORC1, and
2. CDRs generate macropinosomes, which convey extracellular nutrients inside the cells to activate mTORC1.

The critical role of mTORC1 in podocyte homeostasis has been previously reported. We demonstrated that non-diabetic podocyte-specific TSC1 KO mice, in which mTORC1 activity in podocytes is constitutively elevated, displayed severe glomerular dysfunction that shared many similar pathological phenotypes with diabetic nephropathy, including mesangial expansion and podocyte loss/detachment, leading to end-stage renal disease by 14 weeks after birth (Inoki *et al*., 2011). Similarly, genetic ablation of raptor, an essential mTORC1 component in podocytes, causes developmental problems in podocytes and slowly progressive glomerulosclerosis (Godel *et al*., 2011). Moreover, several studies have proposed that mTORC1-mediated podocyte hypertrophy is a protective mechanism that preserves glomerular function (Nishizono *et al*, 2017; Puelles *et al*., 2019; Zschiedrich *et al*, 2017). Thus, the appropriate activity of mTORC1 plays a crucial role in maintaining the physiological functions of podocytes (Fantus *et al*, 2016; Huber *et al*, 2012). The findings reported here suggest that podocyte CDRs have unique cellular functions that control the mTORC1 pathway.

In summary, through a combination of *in vitro, ex vivo*, and *in vivo* experiments, we have identified the expression of CDRs in podocytes and revealed that they function in growth factor signaling. Although the critical roles of mTORC1 in maintaining podocyte viability and function have been well documented, the molecular mechanisms underlying growth factor-induced mTORC1 activity in cells have not been well studied. Our observations suggest that CDRs act as key signaling hubs that trigger growth factor-induced mTOR signaling, including both mTORC2 and mTORC1. The findings we report here should shed light on the biological/biochemical research areas of podocytes and provide additional options to target cellular mTOR activity to control mTOR-related kidney diseases.

## MATERIALS and METHODS

### Reagents, plasmids, and antibodies

Recombinant murine EGF (315-09), murine PDGF-BB (315-18), and murine IFN-γ (315-05) were obtained from Peprotech. EIPA (1154-25-2) was obtained from TOCRIS. MK2206 (A3010) was purchased from APExBio. Wiskostatin was from Abcam (ab141085). Collagen type I solution from rat tail (C3867-1VL) and collagenase (C9263) were purchased from Sigma–Aldrich. Paraformaldehyde (16%, 28908) was from Thermo Fisher Scientific. Glutaric dialdehyde (25%, 16220) was from Electron Microscopy Sciences, Roswell Park Memorial Institute (RPMI). 1640 medium (C11975500BT), Opti-medium (Gibco, 31985-070), low-glucose Dulbecco’s Modified Eagle Medium (DMEM) (C11885500BT), Hank’s balanced salt solution (HBSS) (14025-092), HBSS without magnesium and calcium (14175-079), and fetal bovine serum (10099141C) were obtained from Gibco. Dulbecco’s phosphate-buffered saline (DPBS) (B220KJ) was purchased from BasalMedia. The anti-mycoplasma drug plasmocin prophylactic (ant-app) was purchased from InvivoGen. Penicillin G Na salt (S17030) was purchased from Shanghai YuanyeBio-Technology. Streptomycin sulfate (A610494) was obtained from Sangon Biotech. Lipofectamine 2000 reagent (11668-019) was purchased from Invitrogen. Rhodamine phalloidin (RM02835) was obtained from ABClonal. Mounting Medium with DAPI (ab104139) was from Abcam. The AKT-GFP plasmid (#86637), mSIN1-GFP plasmid (#72907), and Btk-PH-GFP plasmid (#51463) were purchased from Addgene. Cell strainers of 100 (15-1100), 70 (15-1070), and 40 μm (15-1040) were obtained from Biologix. Fluorescein isothiocyanate-labeled dextran with an average molecular weight of 70,000 (FDx70) (D1823) was obtained from Invitrogen. Lucifer yellow (LY) (abs47048137) was obtained from Absin (Shanghai, China). The following materials were used for the immunofluorescence staining assays: WT1 (ABclonal, A17006, 1:50), Rab5a (CST, #21433, 1:50), AKT (CST, #9272 and #2920, 1:50), pAKT (CST, #4060, 1:50), rictor (Proteintech, 27248-1-AP, 1:50), SH3YL1(Abcam, ab154123, 1:50), cortactin (ABclonal, A9518, 1:50), N-WASP (ABclonal, A2576, 1:50), mTOR (CST, #2983, 1:50), nephrin (Abcam, ab216341, 1:50), synaptopodin (Proteintech, 21064-1-AP, 1:50), goat anti-rabbit lgG H&L (Alexa FluorR 488) (abcam, ab150081, 1:500), goat anti-mouse IgG (H+L) Alexa Fluor 546 (Invitrogen, A-11030, 1:500). The western blotting materials were: AKT (CST, #9272, 1:2000), phospho-AKT (Ser473) (CST, #4060, 1:1000), MAPK (ERK1/2) (CST, #4695, 1:2000), phospho-MAPK (ERK1/2) (CST, #4376, 1:1000), S6K (CST, #2708, 1:2000), Phospho-S6K (The389) (CST, #9234, 1:1000), and HRP-conjugated anti-rabbit IgG (GE Healthcare, NA934V, 1:5000).

### Cell culture and inhibitor treatments

The mouse podocyte cell line, MPC5, was obtained from Tongpai Biotechnology (Shanghai, China). The cells were cultured in RPMI 1640 medium containing recombinant murine IFN-γ (50 U/mL), 10% FBS, and 1% penicillin-streptomycin at 33 °C as described previously (Mundel *et al*, 1997b). Differentiation was achieved by culturing the cells for 14 d at 37 °C in RPMI 1640 with 10% FBS and antibiotics. Plasmocin prophylaxis was added at a ratio of 1:1,000 to prevent mycoplasma contamination. All experiments were performed using day 14 podocytes. Inhibitor treatments included EIPA (75 μM), MK2206 (2 μM), and Wiskostatin (10 μM). All inhibitors were applied for 30-min pre-treatments in low-glucose DMEM at the concentrations described above. For amino acid starvation, HBSS (with magnesium and calcium) was added for 30 min after extracting the low-glucose DMEM.

### Scanning electron microscopy

SEM samples of mouse glomeruli were prepared as previously described (Yoshida *et al*., 2021). Briefly, after mice were anesthetized and perfused with PBS followed by fixation buffer (2.5% glutaraldehyde, 2% formaldehyde in 0.1 M cacodylate buffer), the kidneys were dissected, incubated in fixation buffer for 1 h at 25±2 °C, and then stored at 4 °C. Samples were submitted and processed for SEM by the University of Michigan Microscope and Image Analysis Core Facility using standard procedures (Yoshida *et al*., 2021). For advanced SEM observations, we used the Thermo Fisher Helios 650 Nanolab SEM at the Michigan Center for Materials Characterization. SEM samples of the MPC5 cells were prepared as follows:

Cells were prepared on 17-mm collagen-coated coverslips, incubated in low glucose DMEM without FBS for 18 h, stimulated by 16 nM EGF or 2 nM PDGF for 5 min, and then fixed using 2.5% glutaraldehyde (in Sorensen buffer, 35.76 g Na2HPO4 and 9.08 g KH_2_PO_4_ in 1 L ddH_2_O, pH=7.2) at 4 °C overnight. The samples were then dehydrated and embedded in Yimingfuxing Bio. A field emission scanning electron microscope (FEI Apreo S LoVac) was used at the Central Laboratory of Nankai University to examine the CDR structures in the podocytes.

### Immunofluorescence staining and confocal microscopy

After starvation and stimulation, the cells were fixed on 17-mm coverslips with 4% paraformaldehyde in PBS buffer for 30 min and then washed in TBST (Tris 200 mmol/L, NaCl 3 mol/L, pH=7.4). After washing, the cells were permeabilized in 0.1% Triton-X100 for 5 min, blocked in 5% BSA (in TBST) for 30 min, and then washed at 25±2 °C. Primary antibodies were diluted in 5% BSA and cells were incubated in antibody solution at 4 °C overnight. Secondary antibodies were diluted in 5% BSA and the cells were incubated for 2 h at 25±2 °C. Rhodamine phalloidin was diluted as 1:100 in 5% BSA to mark actin, and the cells were incubated for 1 h at 25±2 °C. Coverslips were mounted using a mounting medium with DAPI at 25±2 °C. The samples were examined using a Leica TCS SP5 confocal microscope at the Core Facility of the College of Life Sciences of Nankai University.

### Macropinosome assay *in vitro*

Cells were cultured overnight on coverslips in low-glucose DMEM without FBS. PDGF (2 nM) or EGF (16 nM) were added together with fluorescein dextran (FDx70; 0.5 mg/mL), and the cells were incubated at 37 °C for 5 or 30 min. Cells were fixed with 4% paraformaldehyde (PFA) in PBS at 37 °C for 30 min. The fixed cells were washed three times with DPBS for 10 min at 25±2 °C and mounted. Fifteen phase-contrast and FITC-FDx70 images of each sample were taken using a Live Cell Station (Zeiss Axio Observer Z1) at the Core Facility of the College of Life Sciences of Nankai University. The number of induced CDRs was counted using phase-contrast images and the number of induced macropinosomes was determined by counting FDx70-positive vesicles. More than 5,000 cells were counted from each sample.

### Cell lysates and western blotting

Cells were stimulated with EGF/PDGF (with varied duration), then lysed for 10 min in cold lysis buffer (CHAPS, 40 mM HEPES, 120 mM NaCl, 1 mM EDTA, 10 mM pyrophosphate, 10 mM glycerophosphate, 1.5 mM sodium vanadate, 0.3% CHIPS, pH=7.5). Lysates were centrifuged at 210 *g* for 10 min at 4 °C. The supernatant was mixed with 5x SDS sample buffer and boiled for 5 min. The samples were subjected to SDS-PAGE and western blotting with the indicated antibodies, as described previously.

### CDR assay

The same fixative used for coverslips for immunofluorescence staining was used for the CDR assay. Samples were stained with rhodamine phalloidin and mounting medium with DAPI to count the CDRs and cells. A ZEISS Axio Imager Z1 at the Core Facility of the College of Life Sciences of Nankai University was used to obtain images under a magnification of 20x. Fifteen images were randomly taken from each sample, and more than 2,000 cells were counted per condition. The CDR frequency was calculated as the number of cells showing CDRs in proportion to the total number of cells counted, and the CDR frequency was compared between the inhibitor-treated group and the control group. The average and standard error of the ratios were calculated from at least three independent experiments using a two-tailed Student’s t-test.

### Quantification of signal intensity at CDRs

Images were analyzed using ImageJ. Original images were opened and converted to 8-bit images for quantification. pAKT (or rictor) image was used to generate the THRESHOLD image, which determines the pAKT (or rictor) signal positive area, and saved as “Image 1”. Next, pAKT/AKT (or rictor/AKT) RATIO images were prepared and saved as “Image 2”. “Image 1” and “Image 2” were combined to generate the AND image, which shows the ratio value only on the signal positive area, and this image was saved as “Image 3”. CDR areas on “Image 3” were measured, and the average of the ratio value inside each area was measured.

### Plasmids transfection

The BtK-PH-GFP, AKT-GFP, and mSIN1-GFP plasmids were purified using an EndoFree Midi Plasmid Kit (TIANGEN, #DP108). A plasmid volume of 0.4 μg was applied to each coverslip in the transfection step. A volume of 50 μL Opti-medium with plasmid was added to 50 μL Opti-medium with 2 μL Lipofectamine 2000 Reagent. The mixed Opti-medium was added to 500 μL RPMI 1640 (with FBS, antibiotics, and anti-mycoplasma drugs) for each sample. After 6 h, the medium was replaced with fresh RPMI 1640 medium for 4 h and then changed to low-glucose DMEM for 18 h of starvation. GF treatments, fixation procedures, and observations were carried out as described above in the section “Immunofluorescence staining and confocal microscopy.”

### Isolation of glomeruli

Isolated mouse glomeruli were prepared as previously described (Wang *et al*., 2019). Adult C57BL/6N mice, aged 8-10 wk of age were used in this study. After the mice were anesthetized and killed, the kidneys were carefully removed from each animal, and a few drops of HBSS were applied to avoid drying. None of the HBSS used in the isolation procedure contained any magnesium or calcium. Renal cortices were minced to tiny particles (approximately 1 mm^3^) with a blade and then transferred to 1.5-mL tubes containing 500 μL HBSS. The mixture was centrifuged at 210 *g* for 5 min, after which the particles were transferred to 5 mL collagenase (1 mg/mL in HBSS) to digest for 15 min in a water bath at 37 °C. The resulting liquid was then gently pipetted 20 times at 5-min intervals. The digestion was stopped by adding 5 mL RPMI1640 medium supplemented with 10% FBS, and the mixture was centrifuged at 210 *g* for 5 min. The pellet was resuspended in 5 mL cold HBSS. The resulting mixture was transferred to a 100-μm cell strainer. The large particles were pressed through a 100-μm cell strainer using the rubber plunger of a 1-mL syringe. The filtrate was collected and washed, first through a 70-μm cell strainer and then a 40-μm cell strainer. The fragments that remained on the 40-μm strainer were rinsed with cold HBSS and transferred to a clean cell culture dish. After 2 min of settling, the supernatant was collected and washed through another 40-μm cell strainer. The fragments that remained on this strainer were rinsed with cold HBSS and transferred to another clean cell culture dish. The supernatant was collected and, after 2 min of settling, 0.01% (w/v) polyvinylpyrrolidone (PVP) was added and the mixture was centrifuged at 210 *g* for 5 min to collect the glomeruli.

### SEM and immunofluorescence staining of isolated glomeruli

After isolation, glomeruli SEM samples were directly fixed with 2.5% glutaraldehyde in the Sorensen buffer, as described previously. The samples were sent to Yimingfuxing Bio for dehydration and examined using a field emission scanning electron microscope (FEI Apreo S LoVac) at the Central Laboratory of Nankai University. For immunofluorescence staining of glomeruli, 4% PFA in PBS was used for fixation at 25±2 °C for 1 h. For permeabilization of isolated glomeruli, 0.1% Triton-X100 was used for 10 min with gentle rotation, followed by a 30-min blocking period with 5% BSA. Primary antibodies and rhodamine phalloidin (1:50) were mixed with 5% BSA) for 3 h treatment with gentle rotation. Samples were treated with secondary antibodies (1:200) for 2 h and then DAPI solution (in 5 % BSA, 1:2000) for 10 min. The samples were washed with TBST (5 min X 3) between each step and then mounted in 35% glycerol in TBST. At the end of each step, the mixture was centrifuged (3 min, 210 *g*), and the glomerular pellets were collected. A Leica TCS SP5 confocal microscope was used at the Core Facility of the College of Life Sciences of Nankai University to examine the samples. For each glomerulus, 20-21 images were taken from the top, with a 0.76-μm interval between each image.

### Macropinosome assay *ex vivo*

Isolated glomeruli were treated with LY (0.5 mg/mL) for 5 or 30 mins at 37 °C. After washing three times with PBS with PVP, the glomeruli were fixed in 4% PFA in PBS at 25±2 1)C for 1 h. Actin staining of isolated glomeruli with LY was performed as described in the section above. For each sample, more than 20 glomeruli were counted using a Leica TCS SP5 confocal microscope (Core Facility of the Nankai University College of Life Sciences). For quantification of the LY signal inside glomeruli, more than 40 glomeruli from each group were counted in two independent experiments. The XY-projection images were constructed from confocal scanning images taken from the top 15 μm of each glomerulus and the total LY signals were measured. Actin projection images were used to identify the shape of each glomerulus, and the average LY signal was calculated as (total LY signal per glomerulus)/(area of each glomerulus).

### Statistical analysis

All experiments were conducted at least twice or with at least two biological replicates. Results are expressed as mean ± SEM. P values of less than 0.05 were considered statistically significant. A two-tailed paired Student’s t-test was used for comparisons between two groups (Figure 3H (n=5); Figure 3J (n=5); and Figure S1G (n=5). A two-tailed unpaired Student’s t-test was used for comparisons between the two groups in Figure 4J (n>12), Figure 4L (n>12), and Figure 5G (n>40).

## ACKNOWLEDGEMENTS

This study was supported by Frontiers Science Center for Cell Responses Grants from Nankai University (Grant No. C029205001) and the Shenzhen Science and Technology Program (Grant No. JCYJ20210324120813037). We thank Dr. Joel Swanson from the University of Michigan Medical School for reviewing and editing this manuscript.

## AUTHOR CONTRIBUTIONS

RH initiated, designed, and performed experiments with support from MT, JW, XS, and LW. SY conceived the study, designed the experiments, and wrote the manuscript with support from KI.

## ABBREVIATIONS

CDR: circular dorsal ruffle
PDGF: platelet-derived growth factor
HGF: hepatocyte growth factor
EGF: epidermal growth factor
PI3K: phosphoinositide 3-kinase
mTORC2: mechanistic target-of-rapamycin complex-2
mTORC1: mechanistic target-of-rapamycin complex-1
PIP3: phosphatidylinositol (3,4,5)-triphosphate
PH domain: pleckstrin homology domain
HB-EGF: heparin-binding epidermal growth factor-like growth factor
SEM: scanning electron microscopy
MPC5: mouse podocyte cell line 5
IF: immunofluorescence
FDx70: fluorescein isothiocyanate-dextran 70
LY: lucifer yellow

## SUPPLEMENTARY-FIGURE LEGENDS

**Figure S1.**
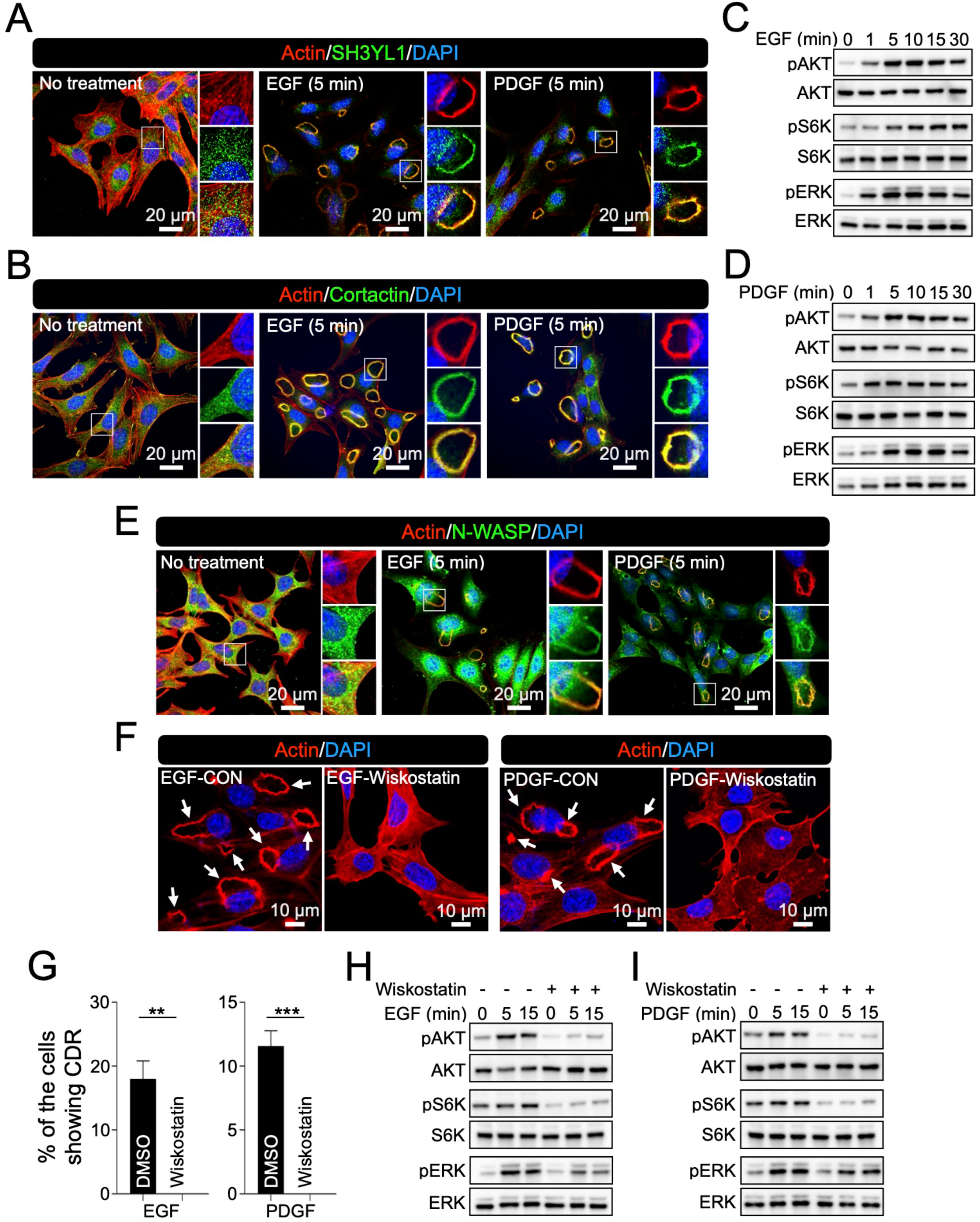
Supplemental data for Figures 2 and 3. (**A, B, and E**) Representative confocal images of actin (red) with SH3YL1 (green in **A**), cortactin (green in **B**), or N-WASP (green in **E**) with and without GF stimulations. Scale bar: 20 μm. (**C and D**) Western blot analysis of AKT, S6K, and ERK signals after EGF (**C**) or PDGF (**D**) stimulation. (**F**) Representative confocal image of MPC5 after GF stimulations with and without N-WASP inhibitor Wiskostatin (shown as Con vs Wiskostatin). Arrows indicate CDRs. Scale bar: 10 μm. (**G**) Results of CDR assays with and without Wiskostatin from five independent experiments. **: p<0.01, ***: p<0.001. Two-tailed paired Student t-test was used for statistics. (**H and I**) Western blot analysis of AKT, S6K, and ERK signals after EGF (**H**) or PDGF (**I**) stimulation with and without Wiskostatin.

**Figure S2.**
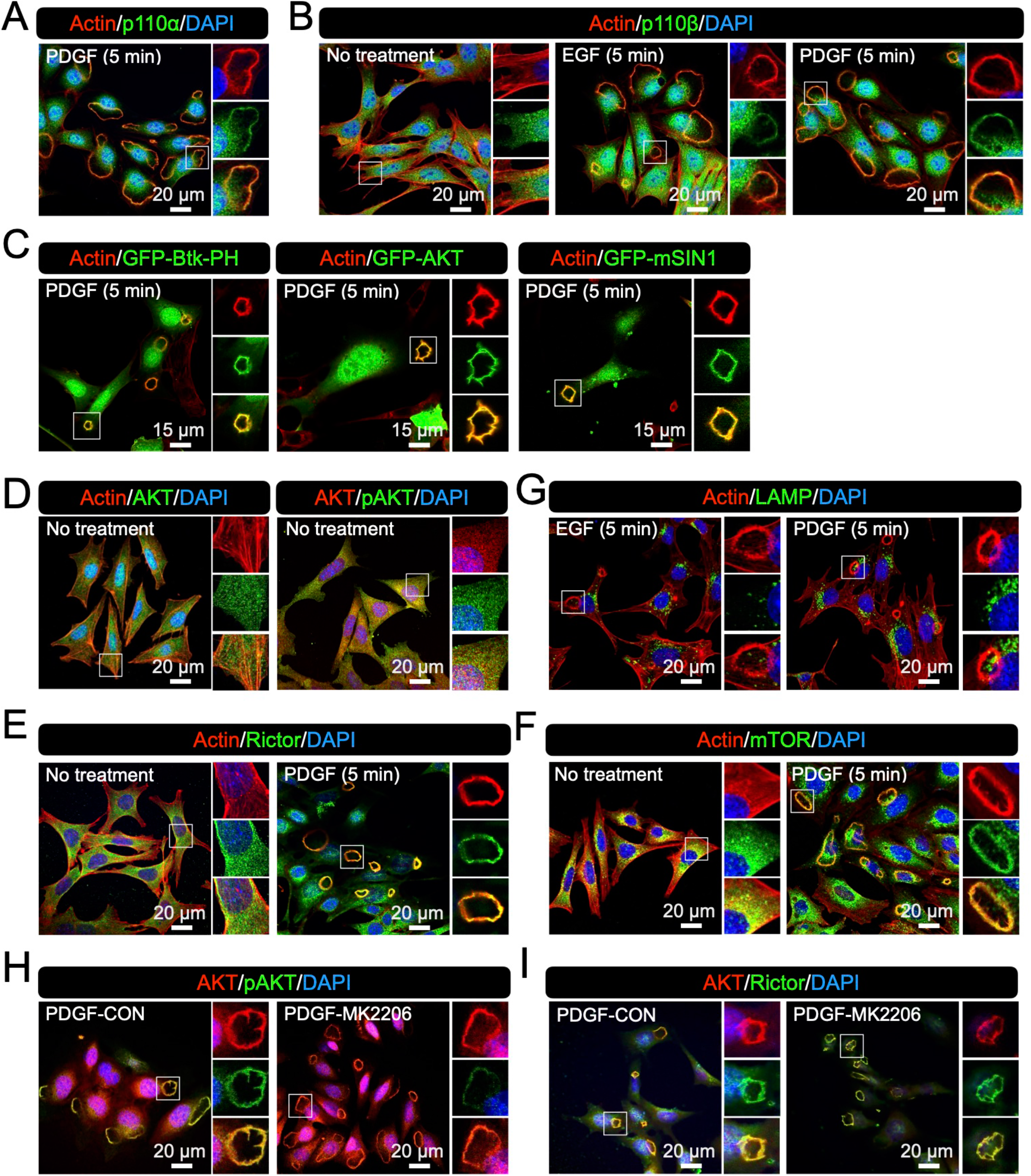
Supplemental data for Figure 4. (**A**) Representative confocal image of actin (red) and p110α (green) in MPC5 after PDGF stimulation. (**B**) Representative confocal image of actin (red) and p110β (green) in MPC5 with and without EGF/PDGF stimulation. (**C**) Representative confocal image of actin (red) with GFP-Btk-PH, GFP-AKT, or GFP-mSIN1 (green) in MPC5 after PDGF stimulation. (**D**) Representative confocal image of actin (red)/AKT (green) and AKT (red)/pAKT (green) in MPC5 with no treatment. (**E and F**) Representative confocal image of actin (red) and rictor (green in **E**)/mTOR (green in **F**) in MPC5 with and without PDGF stimulation. (**G**) Representative confocal image of actin (red) and LAMP (green) in MPC5 after GF stimulations. LAMP, which was not located at the CDRs, was used as a negative control. (**H and I**) Representative confocal image of AKT/pAKT (**H**) and AKT/rictor (**I**) in MPC5 after PDGF stimulation with and without MK2206. Scale bars: 20 μm (**A-B and D-I**), and 15 μm (**C**).

**Figure S3.**
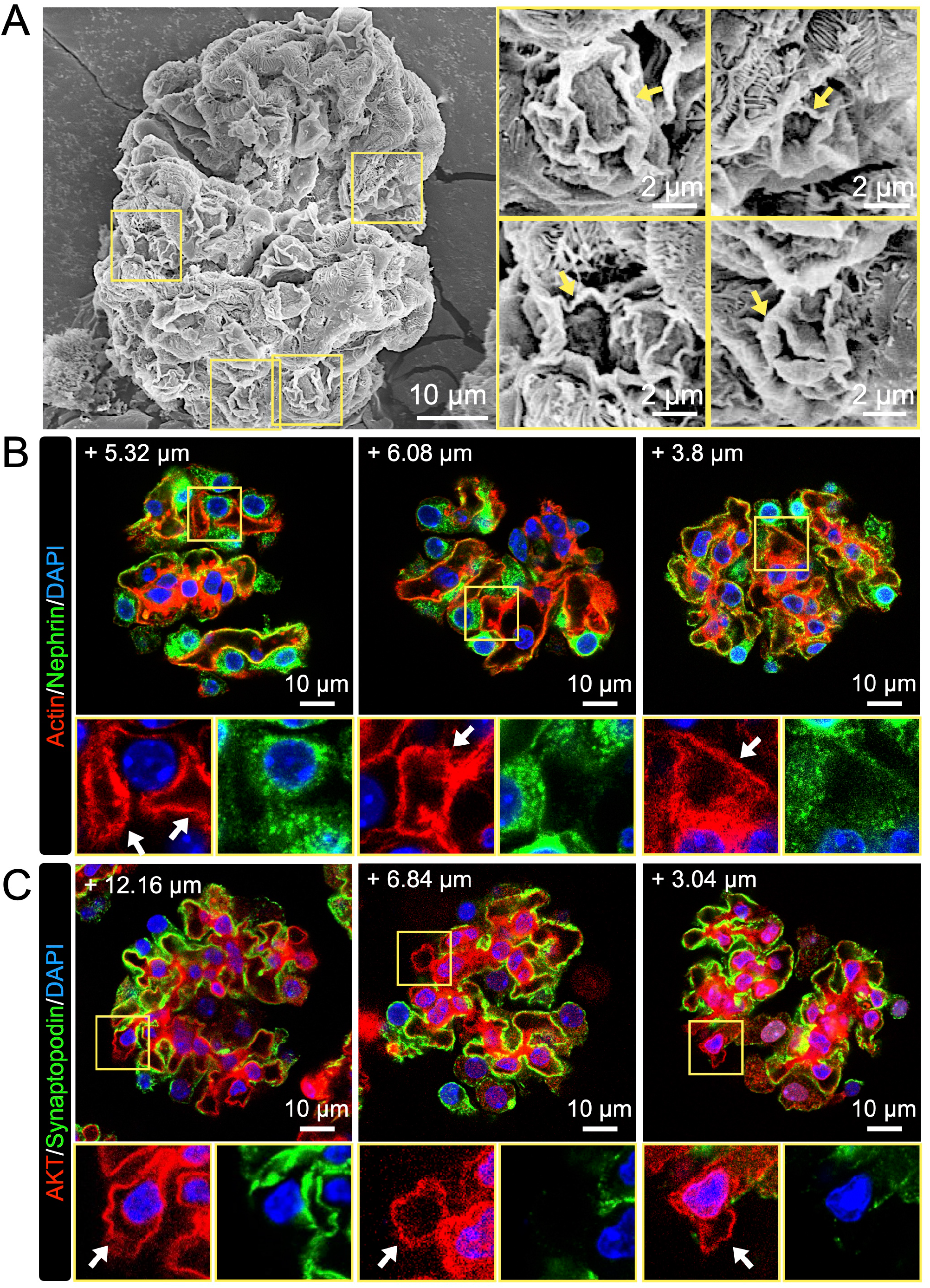
Supplemental data for Figure 5. (**A**) Supplemental high-resolution SEM images of isolated glomerulus showing CDR-like structures. High magnification images of structures resemble morphological steps of CDR formation in MPC5 (arrows). Scale bars: 10 μm and 2 μm (enlarged insets). (**B**) Supplemental confocal images of actin (red) and nephrin (green) in isolated glomeruli. Arrows indicate actin-positive/nephrin-negative structures, presumably CDRs. Three different glomeruli are shown. (**C**) Supplemental confocal images of AKT (red) and synaptopodin (green) in isolated glomeruli. Arrows indicate AKT-positive/synaptopodin-negative structures, suggesting that AKT was recruited to CDRs. Three different glomeruli were shown. The numbers shown in the images indicate the z-axis location of each section (**B and C**). Scale bar: 10 μm (**B and C**).

## REFERENCES

Assady S, Wanner N, Skorecki KL, Huber TB (2017) New Insights into Podocyte Biology in Glomerular Health and Disease. J Am Soc Nephrol 28: 1707–1715

Bar-Sagi D, Feramisco JR (1986) Induction of membrane ruffling and fluid-phase pinocytosis in quiescent fibroblasts by ras proteins. Science 233: 1061–1068

Bernitt E, Dobereiner HG, Gov NS, Yochelis A (2017) Fronts and waves of actin polymerization in a bistability-based mechanism of circular dorsal ruffles. Nat Commun 8: 15863

Berthiaume EP, Medina C, Swanson JA (1995) Molecular size-fractionation during endocytosis in macrophages. J Cell Biol 129: 989–998

Bollee G, Flamant M, Schordan S, Fligny C, Rumpel E, Milon M, Schordan E, Sabaa N, Vandermeersch S, Galaup A et al (2011) Epidermal growth factor receptor promotes glomerular injury and renal failure in rapidly progressive crescentic glomerulonephritis. Nat Med 17: 1242–1250

Chen J, Zeng F, Forrester SJ, Eguchi S, Zhang MZ, Harris RC (2016) Expression and Function of the Epidermal Growth Factor Receptor in Physiology and Disease. Physiol Rev 96: 1025–1069

Chung JJ, Huber TB, Godel M, Jarad G, Hartleben B, Kwoh C, Keil A, Karpitskiy A, Hu J, Huh CJ et al (2015) Albumin-associated free fatty acids induce macropinocytosis in podocytes. J Clin Invest 125: 2307–2316

Cortesio CL, Perrin BJ, Bennin DA, Huttenlocher A, Adams JC (2010) Actin-binding Protein-1 Interacts with WASp-interacting Protein to Regulate Growth Factor-induced Dorsal Ruffle Formation. Molecular Biology of the Cell 21: 186–197

Dharmawardhane S, Schürmann A, Sells MA, Chernoff J, Schmid SL, Bokoch GM (2000) Regulation of macropinocytosis by p21-activated kinase-1. Mol Biol Cell 11: 3341–3352

Egami Y, Taguchi T, Maekawa M, Arai H, Araki N (2014) Small GTPases and phosphoinositides in the regulatory mechanisms of macropinosome formation and maturation. Front Physiol 5: 374

Fantus D, Rogers NM, Grahammer F, Huber TB, Thomson AW (2016) Roles of mTOR complexes in the kidney: implications for renal disease and transplantation. Nat Rev Nephrol 12: 587–609

Fu W, Hall MN (2020) Regulation of mTORC2 Signaling. Genes (Basel) 11

Garg P (2018) A Review of Podocyte Biology. Am J Nephrol 47 Suppl 1: 3–13

Godel M, Hartleben B, Herbach N, Liu S, Zschiedrich S, Lu S, Debreczeni-Mor A, Lindenmeyer MT, Rastaldi MP, Hartleben G et al (2011) Role of mTOR in podocyte function and diabetic nephropathy in humans and mice. J Clin Invest 121: 2197–2209

Gu ZZ, Noss EH, Hsu VW, Brenner MB (2011) Integrins traffic rapidly via circular dorsal ruffles and macropinocytosis during stimulated cell migration. Journal of Cell Biology 193: 61–70

Guo JK, Menke AL, Gubler MC, Clarke AR, Harrison D, Hammes A, Hastie ND, Schedl A (2002) WT1 is a key regulator of podocyte function: reduced expression levels cause crescentic glomerulonephritis and mesangial sclerosis. Hum Mol Genet 11: 651–659

Hasegawa J, Tokuda E, Tenno T, Tsujita K, Sawai H, Hiroaki H, Takenawa T, Itoh T (2011) SH3YL1 regulates dorsal ruffle formation by a novel phosphoinositide-binding domain. J Cell Biol 193: 901–916

Hoon JL, Wong WK, Koh CG (2012) Functions and regulation of circular dorsal ruffles. Mol Cell Biol 32: 4246–4257

Hoxhaj G, Manning BD (2020) The PI3K-AKT network at the interface of oncogenic signalling and cancer metabolism. Nat Rev Cancer 20: 74–88

Huber TB, Edelstein CL, Hartleben B, Inoki K, Jiang M, Koya D, Kume S, Lieberthal W, Pallet N, Quiroga A et al (2012) Emerging role of autophagy in kidney function, diseases and aging. Autophagy 8: 1009–1031

Inoki K, Li Y, Xu T, Guan KL (2003) Rheb GTPase is a direct target of TSC2 GAP activity and regulates mTOR signaling. Genes Dev 17: 1829–1834

Inoki K, Mori H, Wang J, Suzuki T, Hong S, Yoshida S, Blattner SM, Ikenoue T, Ruegg MA, Hall MN et al (2011) mTORC1 activation in podocytes is a critical step in the development of diabetic nephropathy in mice. J Clin Invest 121: 2181–2196

Itoh T, Hasegawa J (2013) Mechanistic insights into the regulation of circular dorsal ruffle formation. J Biochem 153: 21–29

Kay RR (2021) Macropinocytosis: Biology and mechanisms. Cells & Development

Koivusalo M, Welch C, Hayashi H, Scott CC, Kim M, Alexander T, Touret N, Hahn KM, Grinstein S (2010) Amiloride inhibits macropinocytosis by lowering submembranous pH and preventing Rac1 and Cdc42 signaling. J Cell Biol 188: 547–563

Kopp JB, Anders HJ, Susztak K, Podesta MA, Remuzzi G, Hildebrandt F, Romagnani P (2020) Podocytopathies. Nat Rev Dis Primers 6: 68

Lanzetti L, Palamidessi A, Areces L, Scita G, Di Fiore PP (2004) Rab5 is a signalling GTPase involved in actin remodelling by receptor tyrosine kinases. Nature 429: 309–314

Legg JA, Bompard G, Dawson J, Morris HL, Andrew N, Cooper L, Johnston SA, Tramountanis G, Machesky LM (2007) N-WASP involvement in dorsal ruffle formation in mouse embryonic fibroblasts. Mol Biol Cell 18: 678–687

Liu GY, Sabatini DM (2020) mTOR at the nexus of nutrition, growth, ageing and disease. Nat Rev Mol Cell Biol 21: 183–203

Liu L, Doné SC, Khoshnoodi J, Bertorello A, Wartiovaara J, Berggren PO, Tryggvason K (2001) Defective nephrin trafficking caused by missense mutations in the NPHS1 gene: insight into the mechanisms of congenital nephrotic syndrome. Hum Mol Genet 10: 2637–2644

Mendoza MC, Er EE, Blenis J (2011) The Ras-ERK and PI3K-mTOR pathways: cross-talk and compensation. Trends Biochem Sci 36: 320–328

Mundel P, Heid HW, Mundel TM, Krüger M, Reiser J, Kriz W (1997a) Synaptopodin: an actin-associated protein in telencephalic dendrites and renal podocytes. J Cell Biol 139: 193–204

Mundel P, Reiser J, Zúñiga Mejía Borja A, Pavenstädt H, Davidson GR, Kriz W, Zeller R (1997b) Rearrangements of the cytoskeleton and cell contacts induce process formation during differentiation of conditionally immortalized mouse podocyte cell lines. Exp Cell Res 236: 248–258

Nishizono R, Kikuchi M, Wang SQ, Chowdhury M, Nair V, Hartman J, Fukuda A, Wickman L, Hodgin JB, Bitzer M et al (2017) FSGS as an Adaptive Response to Growth-Induced Podocyte Stress. J Am Soc Nephrol 28: 2931–2945

Pacitto R, Gaeta I, Swanson JA, Yoshida S (2017) CXCL12-induced macropinocytosis modulates two distinct pathways to activate mTORC1 in macrophages. J Leukoc Biol 101: 683–692

Palamidessi A, Frittoli E, Garré M, Faretta M, Mione M, Testa I, Diaspro A, Lanzetti L, Scita G, Di Fiore PP (2008) Endocytic Trafficking of Rac Is Required for the Spatial Restriction of Signaling in Cell Migration. Cell 134: 135–147

Palm W, Park Y, Wright K, Pavlova NN, Tuveson DA, Thompson CB (2015) The Utilization of Extracellular Proteins as Nutrients Is Suppressed by mTORC1. Cell 162: 259–270

Puelles VG, van der Wolde JW, Wanner N, Scheppach MW, Cullen-McEwen LA, Bork T, Lindenmeyer MT, Gernhold L, Wong MN, Braun F et al (2019) mTOR-mediated podocyte hypertrophy regulates glomerular integrity in mice and humans. JCI Insight 4

Schell C, Baumhakl L, Salou S, Conzelmann AC, Meyer C, Helmstadter M, Wrede C, Grahammer F, Eimer S, Kerjaschki D et al (2013) N-wasp is required for stabilization of podocyte foot processes. J Am Soc Nephrol 24: 713–721

Stow JL, Hung Y, Wall AA (2020) Macropinocytosis: Insights from immunology and cancer. Curr Opin Cell Biol 65: 131–140

Sun X, Liu Y, Zhou S, Wang L, Wei J, Hua R, Shen Z, Yoshida S (2022) Circular Dorsal Ruffles Disturb the Growth Factor-induced PI3K-AKT Pathway in Hepatocellular Carcinoma Hep3B cells. Cell Commun Signal

Swanson JA (2008) Shaping cups into phagosomes and macropinosomes. Nat Rev Mol Cell Biol 9: 639–649

Swanson JA, Yoshida S (2019) Macropinosomes as units of signal transduction. Philos Trans R Soc Lond B Biol Sci 374: 20180157

Varnai P, Rother KI, Balla T (1999) Phosphatidylinositol 3-kinase-dependent membrane association of the Bruton’s tyrosine kinase pleckstrin homology domain visualized in single living cells. J Biol Chem 274: 10983–10989

Wall AA, Luo L, Hung Y, Tong SJ, Condon ND, Blumenthal A, Sweet MJ, Stow JL (2017) Small GTPase Rab8a-recruited Phosphatidylinositol 3-Kinase gamma Regulates Signaling and Cytokine Outputs from Endosomal Toll-like Receptors. J Biol Chem 292: 4411–4422

Wang H, Sheng J, He H, Chen X, Li J, Tan R, Wang L, Lan HY (2019) A simple and highly purified method for isolation of glomeruli from the mouse kidney. Am J Physiol Renal Physiol 317: F1217–F1223

Ying HZ, Chen Q, Zhang WY, Zhang HH, Ma Y, Zhang SZ, Fang J, Yu CH (2017) PDGF signaling pathway in hepatic fibrosis pathogenesis and therapeutics (Review). Mol Med Rep 16: 7879–7889

Yoshida S, Hoppe AD, Araki N, Swanson JA (2009) Sequential signaling in plasma-membrane domains during macropinosome formation in macrophages. J Cell Sci 122: 3250–3261

Yoshida S, Pacitto R, Inoki K, Swanson J (2018a) Macropinocytosis, mTORC1 and cellular growth control. Cell Mol Life Sci 75: 1227–1239

Yoshida S, Pacitto R, Sesi C, Kotula L, Swanson JA (2018b) Dorsal ruffles enhance activation of Akt by growth factors. J Cell Sci 131

Yoshida S, Pacitto R, Yao Y, Inoki K, Swanson JA (2015) Growth factor signaling to mTORC1 by amino acid-laden macropinosomes. J Cell Biol 211: 159–172

Yoshida S, Wei X, Zhang G, O’Connor CL, Torres M, Zhou Z, Lin L, Menon R, Xu X, Zheng W et al (2021) Endoplasmic reticulum-associated degradation is required for nephrin maturation and kidney glomerular filtration function. J Clin Invest 131

Zdzalik-Bielecka D, Poswiata A, Kozik K, Jastrzebski K, Schink KO, Brewinska-Olchowik M, Piwocka K, Stenmark H, Miaczynska M (2021) The GAS6-AXL signaling pathway triggers actin remodeling that drives membrane ruffling, macropinocytosis, and cancer-cell invasion. Proc Natl Acad Sci U S A 118

Zobel M, Disanza A, Senic-Matuglia F, Franco M, Colaluca IN, Confalonieri S, Bisi S, Barbieri E, Caldieri G, Sigismund S et al (2018) A NUMB-EFA6B-ARF6 recycling route controls apically restricted cell protrusions and mesenchymal motility. J Cell Biol 217: 3161–3182

Zschiedrich S, Bork T, Liang W, Wanner N, Eulenbruch K, Munder S, Hartleben B, Kretz O, Gerber S, Simons M et al (2017) Targeting mTOR Signaling Can Prevent the Progression of FSGS. J Am Soc Nephrol 28: 2144–2157

